# Individual nephron proteomes connect morphology and function in proteinuric kidney disease

**DOI:** 10.1101/194795

**Authors:** Martin Höhne, Christian K. Frese, Florian Grahammer, Claudia Dafinger, Giuliano Ciarimboli, Linus Butt, Julia Binz, Matthias J. Hackl, Mahdieh Rahmatollahi, Martin Kann, Simon Schneider, Mehmet M. Altintas, Bernhard Schermer, Thomas Reinheckel, Heike Göbel, Jochen Reiser, Tobias B. Huber, Rafael Kramann, Tamina Seeger-Nukpezah, Max C. Liebau, Bodo B. Beck, Thomas Benzing, Andreas Beyer, Markus M. Rinschen

**Affiliations:** Department II of Internal Medicine, University of Cologne, Cologne, Germany; Center for Molecular Medicine Cologne (CMMC), University of Cologne, Cologne, Germany; Cologne Excellence Cluster on Cellular Stress Responses in Aging Associated Diseases (CECAD), University of Cologne, Cologne, Germany; Systems Biology of Ageing Cologne (Sybacol), University of Cologne, Cologne, Germany; Department of Medicine III, University Medical Center Hamburg-Eppendorf, Hamburg, Germany; Department of Medicine IV,Medical Center and Faculty of Medicine - University of Freiburg, Freiburg, Germany; Department of Internal Medicine D, Münster, Germany; Rush University Medical Center, Chicago, IL, USA; Institut of Molecular Medicine and Cell Research, Faculty of Medicine, University of Freiburg, Freiburg, Germany; Institute of Pathology, University Hospital Cologne; BIOSS Centre for Biological Signalling Studies and Center for Biological Systems Analysis (ZBSA), Albert-Ludwigs-University, Freiburg, Germany; Division of Nephrology, RWTH Aachen University, Germany; Department I of Internal Medicine, University of Cologne, Cologne, Germany; Department of Pediatrics, Division of Pediatric Nephrology, University Hospital of Cologne, Germany; Department of Human Genetics, University Hospital Cologne, Germany

## Abstract

In diseases of many parenchymatous organs, heterogenous detoriation of individual functional units determines the clinical prognosis. However, the molecular characterization of these subunits remains a technological challenge that needs to be addressed in order to better understand pathological mechanisms. Sclerotic and proteinuric glomerular kidney disease is a frequent and heterogeneous disease which affects a fraction of nephrons, glomeruli and draining tubules, to variable extents, and for which no treatment exists. Here, we developed and applied an antibody-independent methodology to investigate heterogeneity of individual nephron segment proteomes from mice with proteinuric kidney disease. This “one-segment-one-proteome-approach” defines mechanistic connections between upstream (glomerular) and downstream (tubular) nephron segment populations. In single glomeruli from two different mouse models of sclerotic glomerular disease, we identified a coherent protein expression module consisting of extracellular matrix protein deposition (reflecting glomerular sclerosis), glomerular albumin (reflecting proteinuria) and LAMP1, a lysosomal protein. This module was associated with a loss of podocyte marker proteins. In an attempt to target this protein co-expression module, genetic ablation of LAMP1-correlated lysosomal proteases in mice could ameliorate glomerular damage. Furthermore, individual glomeruli from patients with genetic sclerotic and non-sclerotic proteinuric diseases demonstrated increased abundance of lysosomal proteins, in combination with a decreased abundance of the mutated gene products. Therefore, increased glomerular lysosomal load is a conserved key mechanism in proteinuric kidney diseases, and the technology applied here can be implemented to address heterogeneous pathophysiology in a variety of diseases at a sub-biopsy scale

## Introduction

Many organs consist of repetitive functional subunits at the scale of a few micrometers, among these are islets of the pancreas, liver lobules and kidney nephrons. In a variety of diseases, decay in individual unit function and morphology determines organ function and clinical prognosis (for examples, see (1–4)). Since morphologies (as obtained in histology of biopsies) are frequently ambiguous, it has long been hoped that molecular patterns in native biomaterial provide a “molecular diagnosis”, thereby revealing treatment options and personalized prognosis. In fact, in many of these diseases, transcriptomic patterns have been analyzed and yielded first insights into classification and mechanisms of diseases (“integrative genomics”) (5–8). However, transcripts can only partially explain protein abundances, which is why the attention is shifting towards proteome data, especially in its targeted form (9). Unfortunately, only a few, highly specialized analytical pipelines analyzing pathophysiological processes at the functional unit level are developed, and they have not been applied to human disease samples so far, mainly because the mass spectrometer sensitivity is commonly considered to be insufficient (10). Immunostainings are an alternative approach. However, antibodies have a number of other limitations, among these recognition of unspecific epitopes, limited multiplexity and inaccurate quantification of signal intensities caused by background signals, autofluorescence and signal saturation (11, 12). Finally, single cell approaches, although powerful in assessing heterogeneity, disrupt the initial tissue structure and allow only limited insights into extracellular matrix protein abundance, a strong pathological criterion in a variety of fibrotic and chronic diseases. Therefore, antibody-independent mass spectrometry-based proteome analysis at the *functional unit level* would be a valuable alternative to approach patho-mechanisms and molecular diagnosis in human diseases

The human kidney is one of the most complex parenchymatous organs and consists of up to one million functional units, the nephrons. Each nephron consists of a renal glomerulus, the filtration barrier, and the draining tubule. The glomerulus is the site of the filtration barrier which consists of podocytes, endothelial cells and the glomerular basement membranes. The ultrafiltrate is passed to a draining tubule which consists of various segments with defined functions. Chronic kidney disease (CKD) is one of the most severe risk factors for cardiovascular events and stroke, and is characterized by individual nephron function decay (13). Decay in glomerular function is diagnosed frequently as proteinuric kidney disease, such as in focal segmental glomerulosclerosis (FSGS). FSGS is triggered by a variety of causes, including genetic mutations (WT1, NPHS1), and chemical and inflammatory stimuli (14). These insults lead to loss of podocyte function and sclerosis, typically in a heterogenous pattern

Here, we introduce a scalable method to directly extract quantitative proteomic information from *single* units of the kidney - single glomeruli and single tubule segments. With this technology we are able to quantify the proteomes in kidney nephron segments – glomeruli and tubules – consisting of as little as 200 cells (15) and monitor podocyte marker proteins from as little as 80 podocytes per glomerulus independently of antibodies (16). We here develop an analytical pipeline to integrate causative disease mechanisms in heterogeneous nephron populations at a sub-biopsy scale

## Results and discussion

To analyze the proteome of a single glomerulus, we modified a protocol which was designed for very small sample amounts (17). Commonly used C18 based sample preparation protocols utilize relatively large volumes and has a sub-optimal sample recovery rate. In contrast, this protocol minimizes sample loss by tight binding of proteins and peptides to carboxylated magnetic beads during protein purification (Fig. S1A), making it compatible with most comprehensive lysis buffers (up to 10% SDS). We microdissected wildtype mouse glomeruli and subjected them – one at a time – to proteomic analysis (Fig. S1B). The ultrasensitive sample preparation method largely outperformed the C18-based, “standard” sample preparation (stagetips, Fig. S1C). Proteomic analysis by LC-MS/MS showed a substantial ion current signal from a single mouse glomerulus (Fig. 1A). Expectedly the method was even more successful for larger human glomeruli (Fig. 1B). The single glomerulus datasets contained core podocyte proteins such as nephrin, ACTN4, podocin and CD2AP, proteins of the basement membrane such as collagen type 4 and markers of mesangial cells such as desmin. Next, we analyzed the proteome of anatomically defined, microdissected mouse tubule segments, which is to our knowledge the first proteomic analysis of these structures that complements recent transcriptome acquisitions (18). Again, shotgun proteomic analysis could clearly distinguish single proximal tubules (S1 segments), thick ascending limbs and cortical collecting ducts by means of known marker protein expression (Fig. 1C) and on global proteome level (Fig. S2A,B). Expectedly, thicker segments yielded more peptides than thinner segments, such as thick acending loop of Henle (Fig. S2C). In proximal tubules, the data covered more than 1500 proteins, and the abundance spanned 4 orders of magnitude in single tubules (Fig. S2D). In a similar fashion, we also determined the proteome of single proximal tubules from humans (Fig. S2E-F)

**Figure 1:**
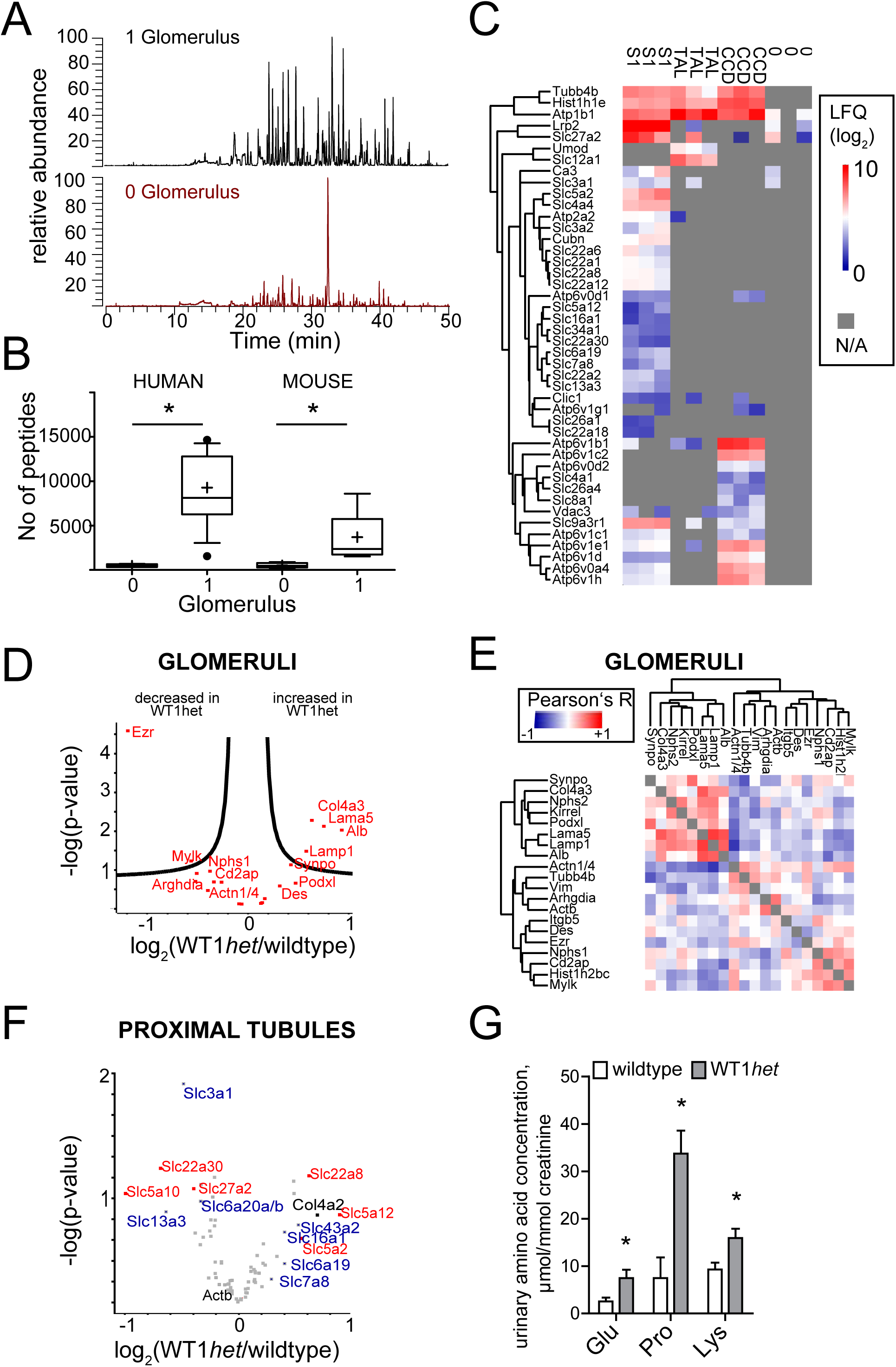
Development of a single segment proteomic method and application to renal segments of the WT1 heterozygous mouse. A. Total ion current during LC-MS/MS acquisition of samples from 0 (vehicle solution only) and 1 mouse glomerulus. B. Comparison of identified peptides from mouse and human glomeruli as determined by ultrasensitive proteomics and MaxQuant analysis (n = 12 single glomeruli from 3 mice and n = 18 single glomeruli from three humans). + = mean, - = median, * = p<0.01 in a two-tailed t-test. Boxes indicate 25-75% percentile. Outliers beyond the 95% percentile are marked with dots. C. Analysis of single, microdissected tubules from mouse kidney cortex. S1 proximal tubules, cortical thick-ascending limb (TAL) and cortical collecting ducts (CCDs) are clearly discernible by proteomic analysis as compared to “empty” (0) samples. A selection of proteins is clustered with their gene symbol. The proteins in the first three rows are non-segment specific tubular proteins such as Na/K ATPase (Atp1b1). D. Volcano plot of single glomeruli. Proteomics analysis of glomeruli obtained from a WT1 heterozygous KO mouse (Wt1het) and wildtype mice as controls. Negative log(p-value) of a two-tailed t-test is plotted against a log2 fold change of Wt1het/wildtype. Quantification is based on n=20 single glomeruli from n=3 different Wt1het animals, and n=20 single glomeruli from n=3 different wildtype controls. E. Hierarchical clustering of correlation coefficients across samples. F. Volcano plot quantification of single S1 tubules microdissected from wildtype vs control mice. Amino acid transporters are marked with blue, and other transporters are indicated in red. G. Selective aminoaciduria of proteinuric *Wt1het* mice for proline (a substrate of SLC6A20), and lysine (a substrate of SLC3A1) and glutamate (a lower-affinity substrate of SLC13A3). n=6, * p<0.05 two-tailed t-test

Since proteomes from single functional units could be resolved, we studied a disease model of WT1 haploinsufficiency. WT1 is a podocyte-specific transcription factor controlling expression of various podocyte-specific genes, *Wt1het* mice develop proteinuria and partially sclerotic lesions at the age of 14 weeks resembling FSGS (19). Shotgun proteomics applications, as utilized above for initial testing of the method, are inherently limited due to stochastic acquisition and undersampling (20).To address heterogeneity and protein expression wiring independently of stochastic acquisition, we set up a targeted proteomics assay, specifically parallel reaction monitoring (PRM). We developed a “podocyte function sentinel assay” which comprises 20 proteins (41 peptides), that were selected based on the known, genetically verified importance for podocyte function, and which were all detected in the global glomerular proteomes (Table S1). We quantified these proteins using the assay in single glomeruli from mice heterozygous for the transcription factor WT1 (*Wt1het*) and wildtype mice (n=3 mice per group, n=7 single glomeruli/mouse) (Table S2). We found that albumin, as well as the extracellular matrix (ECM) proteins collagen type IV and laminin were significantly increased in single glomeruli from *Wt1het* mice (Fig. 1D), a finding consistent with FSGS and proteinuria in these animals (Fig. S3 A,B). Yet, a large variation between different glomeruli was observed (Fig. S3 C illustrates some of the individual measurements for proteins). To illustrate this heterogeneity, we calculated the correlation of all proteins across all glomeruli (correlations with each other), and performed hierarchical clustering of the correlation coefficients. As an example, a strong correlation also occurred between laminin 5 and the lysosomal marker LAMP1 (Fig. 1E). LAMP1 and ACTN1/4 correlated negatively (Fig. 1E). For some proteins, there was no correlation i.e. between nephrin and ACTN1/4 (Fig. 1E)

Next, we analyzed the proximal tubule of the same proteinuric mice. Proximal tubules alter their physiological function due to proteinuria and overwhelming albumin uptake (21). To assess this systematically, we again developed a proteomics sentinel assay to survey proximal tubule status. In total, we monitored 74 proteins that are determining key physiological proximal tubule function (such as metabolite transporters, the Na^+^/K^+^-ATPase and key signaling molecules, Table S3). We isolated proximal tubules from the same *Wt1het* knockout mice as described above and applied the tubule sentinel PRM assay (n=3 mice per group, n= 7 tubules/mouse). In isolated S1 tubular segments, initial correlational analysis revealed distinct clusters of proteins highly associated with each other. Overall regulation of proteins was not determined by known mRNA expression patterns of the different proximal tubule subtypes (S1, S2 or S3, based on (18); Fig. S4A). One of the key tasks of the proximal tubule is the reabsorption of metabolites, such as amino acids, and disruption of this function results in aminoaciduria, which is occasionally observed in nephrotic patients (22, 23). Quantitative analysis (*Wt1het* vs wildtype) revealed that the amino acid carriers SLC3A1, SLC13A3 and SLC6A20a/b were decreased in single *Wt1het* mouse proximal tubules (Fig. 1F). Interestingly, only a few tubules regulated these transporters: those particular tubules that expressed low abundance of these transporters contained high expression of other amino acid transporters with overlapping amino acid transport spectrum (Fig. S4A). Furthermore, these tubules also showed increased abundance of albumin and collagen type 4 (Fig. 1F, Fig. S4A). Consistent with the stochastic decrease in these three transporters, we found moderately increased levels of urinary lysine (one of the substrates of SLC3A1 (24)) and proline (one of the substrates of SLC6A20 (25)), and glutamate (one of the low-affinity substrates of SLC13A3 (26)) in the urine of proteinuric mice (Fig. 1G) – changes in the other amino acids were not significant (Fig. S4B). These data demonstrate that single-segment proteomics can reflect pathomechanisms in proteinuric kidney disease

We decided to follow up on the glomerular function in a second FSGS model using single glomerular proteomics analysis and the same sentinel assay. We used the model of doxorubicin (trade name: Adriamycin) induced FSGS and proteinuria (Fig. S5). We found that in contrast to the *Wt1het* FSGS model, there was a significant decrease in nephrin (gene symbol *Nphs1*) in these glomeruli (Fig. 2A, Table S4). To determine similarities of the *Wt1het* and the doxorubicin damage models, we compared fold changes of the observed proteins between both datasets, unraveling that albumin, laminin, collagen and LAMP1 were upregulated in both models, and ACTN1/4 and CD2AP were decreased in both models (Fig. 2B). We plotted correlation coefficients between all protein pairs in both the doxorubicin model and the *Wt1het* model (Fig. 2C). Interestingly, a common finding between both models was the positive correlation of the lysosomal marker LAMP1 with glomerular albumin and ECM proteins, both markers of glomerular damage (Fig. 2C, upper right quadrant). Thus, for a number of proteins, co-expression is wired across single glomeruli and across damage, and LAMP1 expression was connected with extracellular matrix and albumin amount in single glomeruli in two models, suggesting that an increase in LAMP1 is a common hallmark of the glomerular kidney diseases examined

**Figure 2:**
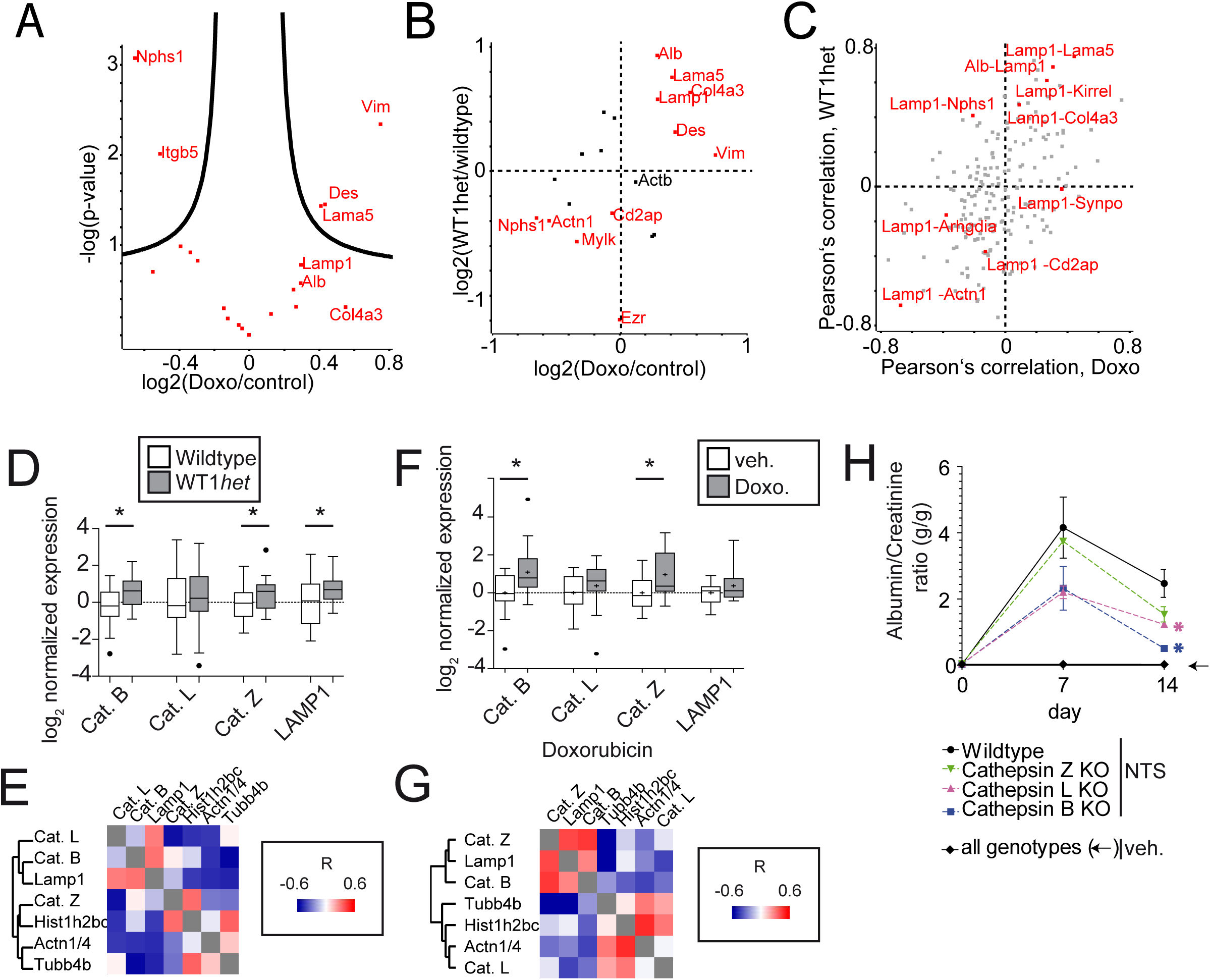
Individual glomeruli analysis reveals a disease-driving module of protein expression in glomerular disease. A. Volcano plot of single glomeruli proteomics analysis of glomeruli obtained from doxorubicin treated and wildtype mice as controls. Negative log(p-value) of a two-tailed t-test is plotted against a log2 fold change of doxorubicin/vehicle. Quantification is based on n=19 single glomeruli from n=3 different animals treated with doxorubicin, and n=19 single glomeruli from n=3 different animals treated with vehicle. B. Correlational analysis of proteins across samples. B. Comparison of log2 fold changes between doxorubicin and Wt1het podocyte damage model. Proteins of interest are marked in red with their gene symbol. The housekeeping protein actin (Actb, depicted in black) remains unchanged in both models. C. Comparison of Pearson’s coefficient between doxorubicin and Wt1het podocyte damage model. Each dot is a correlation pair. D. Fold change comparison of normalized cathepsin B, L, Z and LAMP1 expression in single glomeruli from *Wt1het* and wildtype animals as determined by parallel reaction monitoring. n=16 (control) vs n= 18 (*Wt1het*) glomeruli from 3 different animals. *p<0.05 in a two-tailed t-test. Boxes indicate 25- 75%; and whiskers indicate the 97.5% interval. Median is indicated by a line, mean is indicated by a +. Outliers are marked as dots. E. Correlation analysis of cathepsin B, L, Z and LAMP1 over all single glomeruli from *Wt1het* and wildtype animals. Pearson’s R is clustered according to maximum distance. F. Fold change comparison of cathepsin B, L, Z and LAMP1 in single glomeruli from vehicle and doxorubicin treated animals. n=17 (vehicle) vs n=18 (doxorubicin). G. Correlation analysis of cathepsin B, L, Z and LAMP1 in single glomeruli from vehicle and doxorubicin treated mice. H. Proteinuria (expressed as albumin-creatinine ratio) of cathepsin B, L and Z knockout mice with and without glomerular injury mediated by nephrotoxic serum (NTS). *p<0.05 in a one-way ANOVA followed by Tukey’s post test)

LAMP1 is a structural protein of the lysosome, and may thus not be suitable as a key therapeutic target in renal disease. Therefore, we focused on components of the lysosomes of higher accessibility which could be better druggable targets. Cathepsin L but also other cathepsin proteases are expressed in cultured podocytes and possibly functional in podocytes based on degradomics data (27–30). Since cathepsins can also act outside of lysosomes (31), we investigated whether lysosomal (LAMP1) abundance is linked to cathepsin abundance. To this end, we designed a targeted proteomic assay to monitor the expression of the three proteases (cathepsin B, L, and Z) and LAMP1 (Table S5). In single glomeruli of the *Wt1het* model of podocyte damage, there was a significant increase of cathepsin B and Z, while cathepsin L was increased, but not significantly (Fig. 2D). Both cathepsin B and cathepsin L correlated significantly (p<0.0001) with LAMP1 abundance (Fig. 2E). Somewhat similar, in single glomeruli from proteinuric doxorubicin-treated mice, cathepsin B and Z were found to be increased (Fig. 2F), and both cathepsin B and Z significantly correlated with LAMP1 abundance (Fig. 2G). To systematically test if LAMP1- correlated cathepsins may be causative for glomerular damage, we used constitutive knockout mice for cathepsin B, L and Z. Under baseline conditions, the knockout animals were not proteinuric (Fig. S6A). However, all three knockout strains, especially cathepsin B knockouts, showed more resistance and faster recovery after glomerular damage upon podocyte injury induced by nephrotoxic serum (Fig. 2H, Fig. S6B). These findings demonstrate that single unit proteomic analysis of glomeruli can resolve orchestrated glomerular cathepsin activity, thereby adding LAMP1-correlated cathepsin B as an important causal mediator in different modes of glomerular injury

We found that in two independent models of glomerular disease, LAMP1 was part of a protein co-expression module consisting also of cathepsin proteases, albumin and ECM proteins. Therefore, we asked if this module of proteins could also be found to be disease driving in human proteinuric kidney diseases. We compared the proteome of single glomeruli from patients with nephrotic syndrome with those from tumor nephrectomy samples from adult patients, and again measured the proteome of each single one separately. In single glomeruli from a nephrectomy sample from a five year old patient with primary idiopathic steroid-resistant FSGS, abundance of LAMP1 and SCARB2, an alternative lysosomal marker protein was significantly increased (Fig. 3A). In addition, increased abundance of extracellular matrix proteins was detected compared to a control kidney (Fig. 3A). We also microdissected and measured single glomeruli from nephrectomy samples from two patients with congenital nephrotic syndrome and MCD caused by nephrin (*NPHS1*) mutation. (Table S6 for the dataset, Table S7 for details on the clinical presentation, and Fig. S7 for histology of patients and controls). We identified more than 2000 proteins in this dataset and quantified more than 1000 proteins in all samples. There was a strong reduction of nephrin (NPHS1) in the samples obtained from the glomeruli with *NPHS1* mutation (Fig. 3B), and lysosomal marker LAMP1 was increased. These results could be confirmed when comparing single glomeruli of the second *NPHS1* patient with those of a second control kidney (Fig. 3C). As an ancillary finding, we found reduced amounts of mitochondria in all three datasets analyzed as compared to control cells (Fig. S8). Furthermore, the mass spectrometry data was consistent with the histopathological diagnosis of an enlarged mesangial matrix in one patient (Fig. 3D), which translated into an increased fraction of collagen proteins in the single glomerular proteomes (Fig. 3D). To see if we could reduce the amount of tissue even more, we used laser dissection microscopy from 10µm thick cryosections. We could obtain clear glomerular or tubular proteome patterns which included important mediators of podocyte signaling by proteomic analysis (Fig. S9)

**Figure 3:**
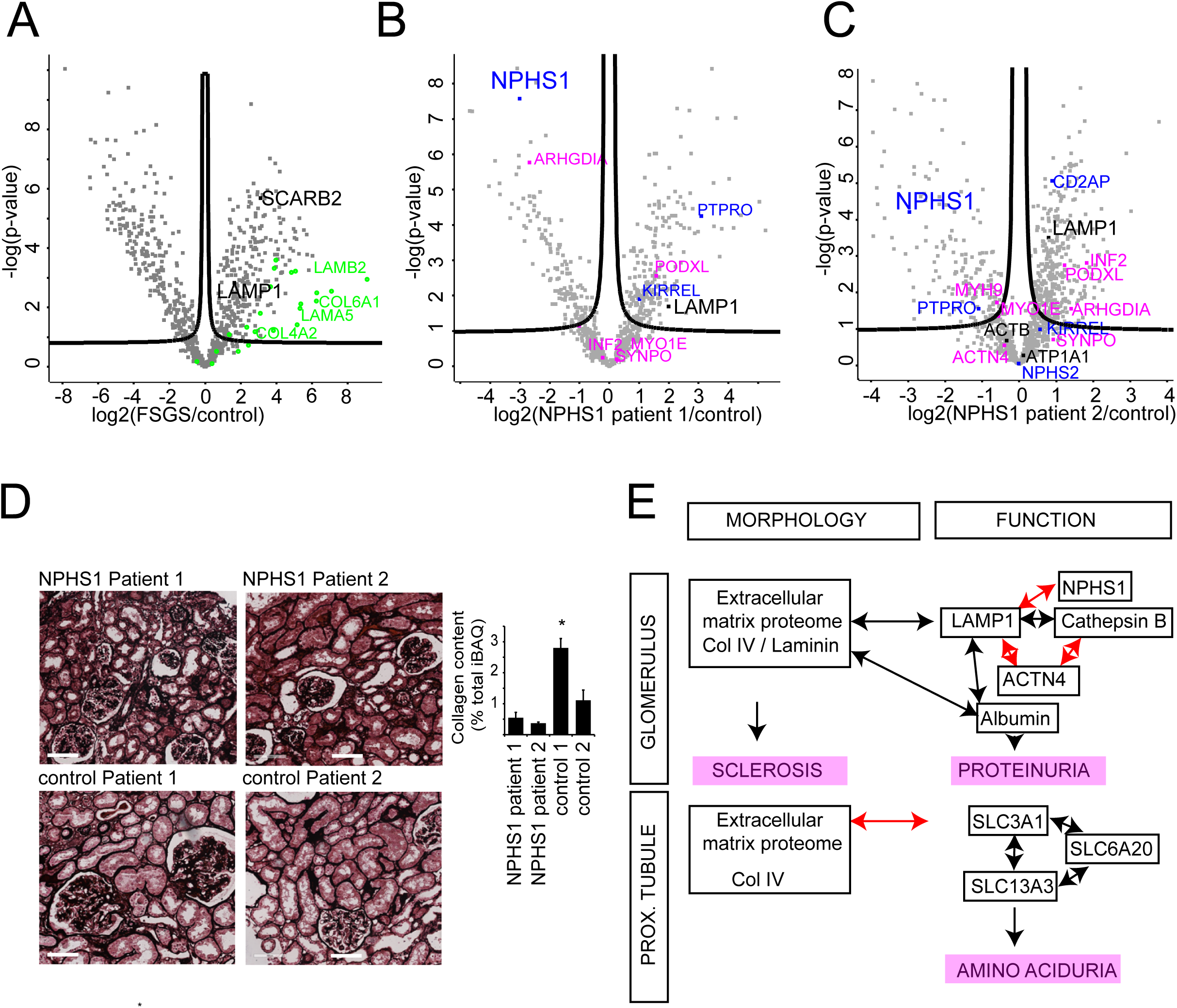
Application of ultrasensitive proteomics on patient material identifies causative mechanisms and increased lysosomal abundance. A. Application of ultrasensitive proteome analysis to single glomeruli from a five year old male patient with primary and idiopathic FSGS; and comparison to a control kidney. Extracellular matrix proteins (green) are increased in glomeruli from FSGS. LAMP1 and SCARB2, two lysosomal markers (marked in black), are increased in FSGS. N = 6 glomeruli/patient. All test are two-tailed t-tests, with the significance of the test (-log10) plotted against the log2 fold change of disease/control. The curved line determine significance after correction for multiple testing (FDR = 0.05. s0 = 0.1). B. Application of ultrasensitive proteome analysis to single glomeruli from one patient (NPHS1 patient 1, n = 7 single glomeruli per patient) with congenital nephrotic syndrome and *NPHS1* mutation; and comparison to a control kidney (control 1, n = 7 single glomeruli). Proteins known to cause genetic glomerular disease are labeled with their gene symbols (pink: actin cytoskeletal proteins, blue: slit diaphragm proteins, and black: other proteins of interest). C. Application of ultrasensitive proteome analysis to single glomeruli from a patient (patient 2) with congenital nephrotic syndrome, minimal-change disease and NPHS1 mutation; and comparison to a control kidney. N=8 glomeruli. Color coding as in Panel B. D. Methenamine-Silver staining of the four patients analyzed in this study, and fraction of iBAQ belonging to collagens in the corresponding datasets. *p<0.05 in a one-way ANOVA with Tukey’s post-test. E. Summary of discovered protein modules discovered by individual nephron proteomics to connect morphology and function. Double-headed arrows indicate correlation of protein expression. Black = positive correlation, red = negative correlation

To summarize, we successfully applied advanced sample preparation (32) to analyze protein composition of kidney tissue at a single-unit level resolution. The “one-glomerulus- one-proteome” approach described herein allows studying *correlated variation* of molecular markers across nephrons and within a kidney, and to functionally link them to both morphology and physiology (Fig. 3E). An LMD-coupled proteomics approach to glomerular disease has shown great success for identifying high abundant extracellular proteins and thereby identifying novel disease mechanisms (33). By improving sensitivity, and utilizing targeted proteomics, we here obtain molecular information also on lower-abundant intracellular signaling proteins from individual glomeruli, which allows definition of correlative modules. These can be targeted therapeutically, and we identify cathepsin B, LAMP1, extracellular matrix proteins and albumin as key components of a disease-driving protein module. Historically, our understanding of renal physiology is derived from the functional analysis of rodent single nephron segments such as kidney tubules (34) or glomeruli (35). Here, we utilize these techniques to allow assessment of physiological and molecular variability in diseases. By converting tissue at the sub-biopsy level into computable datasets, single unit proteomics can uncover disease-driving protein modules which connect morphology and function across segments and individuals. Leveraging intraindividual variability of protein expression within functional units can serve as a blueprint to understand pathobiology also in other organs with similar-sized repetitive functional units

## Acknowledgements

We acknowledge the patients and their parents who donated biomaterial for this study. M.M.R. was supported by a UOC Postdoc grant (DFG Exzellenzinitiative). This study was supported by the DFG (CRC 1140 KIDGEM to FG and TBH, CRC 992 and HU 1016/8-1 to TBH). TBH was supported by the BMBF (01GM1518C), by the European Research Council (ERC grant 616891 and H2020-IMI2 consortium BEAt-DKD) and by the Excellence Initiative of the German Federal and State Governments (BIOSS, EXC-294). M.C.L. and B.S. are supported by the BMBF (01GM1515E). A.B. was supported by the BMBF (SyBACol, FKZ0315893A). We thank Jeroen Krijgsveld (Heidelberg) for donating a custom-made strong-field magnet. The authors thank Ruth Herzog, Ursula Cullmann, Astrid Wilbrand- Hennes and Jan Krüger for excellent technical support

## Authors contributions

M.H., C.F., F.G., J.B., C.D., H.G., S.S., M.M.R., performed experiments. C.F., G.C., M.J.H.,B.S., M.C.L, T.S-N., R.K., J.R., M.M.A., T.R., M.R., T.B.H., R.K., T.S-N. contributed new tools, reagents, animal models or biomaterial. F.G., B.B.B., M.M.R. analyzed data. M.H., F.G. T.B.H., T.B., A.B., M.M.R. interpreted results. M.M.R. conceived and designed study.M.H. and M.M.R. wrote the paper with input from all coauthors

## Methods

### Statistics

Statistics were performed as indicated in the figure legends or in the “bioinformatics section” of the accompanying paper

### Study approval

All investigations were conducted in accordance with the principles of the Declaration of Helsinki. All investigations were conducted after obtaining informed consent from the patients or their parents. All procedures were approved by local ethics committees in Aachen and Cologne. More details on the methods, including animals, proteomics acquisition and data analysis are in the supplementary methods

